# Planar Amorphous Silicon Carbide Microelectrode Arrays for Chronic Recording in Rat Motor Cortex

**DOI:** 10.1101/2023.11.02.565338

**Authors:** Justin R. Abbott, Eleanor N. Jeakle, Pegah Haghighi, Joshua O. Usoro, Brandon S. Sturgill, Yupeng Wu, Negar Geramifard, Rahul Radhakrishna, Sourav Patnaik, Shido Nakajima, Jordan Hess, Yusef Mehmood, Veda Devata, Gayathri Vijayakumar, Armaan Sood, Teresa Thuc Doan Thai, Komal Dogra, Ana G. Hernandez-Reynoso, Joseph J. Pancrazio, Stuart F. Cogan

## Abstract

Chronic implantation of intracortical microelectrode arrays (MEAs) capable of recording from individual neurons can be used for the development of brain-machine interfaces. However, these devices show reduced recording capabilities under chronic conditions due, at least in part, to the brain’s foreign body response. This creates a need for MEAs that can minimize the foreign body response to enable long-term recording. A potential approach to reduce the foreign body response is the use of ultrathin MEAs. Here, we fabricated ultrathin (cross-sectional area: 160 µm^2^) amorphous silicon carbide (a-SiC) MEAs with sixteen electrode channels and implanted them into the motor cortex of seven female Sprague-Dawley rats. A-SiC was chosen as the fabrication base for its high chemical stability, good electrical insulation properties, and amenability to thin film fabrication techniques. Electrochemical analysis and neural recordings were performed weekly for 4 months. MEAs were characterized *in vitro* pre-implantation and *in vivo* using electrochemical impedance spectroscopy and cyclic voltammetry at 50 mV/s and 50,000 mV/s. Neural recordings were analyzed for single unit activity. At the end of the study, animals were sacrificed for immunohistochemistry analysis. We observed statistically significant, but small, increases in 1 and 30 kHz impedance values and 50,000 mV/s charge storage capacity over the 16-week implantation period. Slow sweep 50 mV/s CV and 1 Hz impedance did not significantly change over time. Impedance values increased from 11.6 MΩ to 13.5 MΩ at 1 Hz, 1.2 MΩ to 2.9 MΩ at 1 kHz, and 0.11 MΩ to 0.13 MΩ at 30 kHz over 16 weeks. The median charge storage capacity of the implanted electrodes at 50 mV/s were 58.1 mC/cm^2^ on week 1 and 55.9 mC/cm^2^ on week 16, and at 50,000 mV/s were 4.27 mC/cm^2^ on week 1 and 5.93 mC/cm^2^ on week 16. Devices were able to record neural activity from 92% of all active channels at the beginning of the study, At the study endpoint, a-SiC devices were still recording single-unit activity on 51% of electrochemically active electrode channels. In addition, we observed that the signal-to-noise ratio experienced a small decline of only −0.19 per week. We also classified the units as fast and slow spiking based on the trough-to-peak time. Although the overall presence of single units declined, fast and slow spiking units declined at a similar rate. Furthermore, immunohistochemistry showed minimal foreign body response to the a-SiC devices, as highlighted by statistically insignificant differences in activated glial cells between implanted brains slices and contralateral sham slices, as evidenced by GFAP staining. NeuN staining revealed the presence of neural cell bodies close to the implantation site, again statistically not different from a contralateral sham slice. These results support the use of ultrathin a-SiC MEAs for long-term implantation and use in brain-machine interfaces.

## Introduction

Microelectrode arrays (MEAs) are critical components of neural interfaces that provide an electrical connection between devices such as vision or motor prostheses and neurons, and provide insight into the functional circuitry comprising the complex neuronal networks found within the brain [1]–[6]. Two commonly used cortical MEA devices, the Utah array and Michigan probe, are fabricated using a rigid silicon-based platform [7], [8]. Although these devices have shown the ability to record or stimulate large numbers of neurons in several animal models, and there has been success in human implantation [9], [10], consistent chronic robustness remains a challenge [11]–[14]. This may be due to many factors including insulating material degradation, electrode site delamination, and physical breakage of the devices [14], [15]. However, the reactive tissue foreign body response (FBR) that arises after the invasive probes have been implanted into the cortex remains a major concern [16]–[18]. FBR is critical to minimize as extracellular action potential amplitudes observed at recording sites on penetrating electrode arrays are reduced by tissue encapsulation [19]. Prior work has demonstrated that the FBR can encapsulate arrays in fibrous tissue when implanted in rat motor cortex after a 32-week study [20]. Additionally, as the devices are implanted into the cortex, they rupture blood vessels and may damage neurons and oligodendrocyte support cells [21], [22]. The FBR can be observed to begin within minutes of implantation and prevents the recovery of damaged tissue [21]. As the response continues, the device is encapsulated in a fibrous tissue that damages or appears to push healthy neurons away from the implanted electrodes, reducing the ability for neural recording [23]–[25]. One approach to mitigating this response is to make the cross-sectional dimensions of the penetrating structures, typically the shanks of the MEAs, smaller to minimize trauma associated with surgical implantation, while maintaining the stiffness necessary to penetrate the pial surface and insert into the brain parenchyma without external mechanical support. Structures consisting of features of cross sectional dimensions of only 20 µm^2^ have been shown to trigger reduced FBR [26]. Neuronal cell bodies, confirmed by immunohistochemistry four weeks after implantation, were also observed adjacent to the 20 µm^2^ structures [26]. Sub-cellular scale carbon fiber microwires have also been shown to have reduced FBR when examining implantation sites after 12 weeks *in vivo* [27]. Intrinsically stiff, but highly flexible carbon fiber electrodes with cross sectional areas of 60 µm^2^ were shown to reduce gliosis when compared to commercially available electrode arrays [28]. Additionally, flexible electrode arrays with cross sectional areas of 50 µm^2^ and 10 µm^2^ implanted into cortex through the use of a temporary mechanical shuttle showed excellent recording capabilities in a chronic preparation [29].

In the present study, we investigated amorphous silicon carbide (a-SiC) as both the insulating material and the fabrication base of an ultrathin microelectrode array. A-SiC was chosen due to its relative chemical inertness, good electrical insulating characteristics, and comparatively high stiffness, sufficient to allow designs of cortical penetrating shanks even with ultrathin geometries [30]–[33]. Previous work has demonstrated the low dissolution rate of a-SiC in a saline soak test at 87⁰C for a period of 340 days [34]. A-SiC has been used in stents for patients with acute coronary syndrome [35]. Prior work has also demonstrated the ability of a-SiC arrays to be implanted into rat cortex without an insertion guide, a supporting coating such as polyethylene glycol, or an implant shuttle [32]. Bundled multi-shank a-SiC devices with cross-sectional areas of 60 µm^2^ were found to have well resolved single unit neural recordings in Zebra Finch cortex [36].

For MEA applications, electrode impedance must be low enough to allow neural recording and charge-injection capacity high enough to allow neural stimulation. Sputtered iridium oxide coated a-SiC MEAs have been shown to record neuronal action potentials (single units) from the cortex in a sub-chronic setting [36] and to record stable local field potentials for 16 weeks following implantation [37]. In this study, we assessed the ability of these MEAs to record single units for 16 weeks following implantation. This was done by identifying the presence of single units in neural recordings and calculating parameters such as the signal-to-noise ratio of recordings [38]. We also assessed the prevalence of fast-spiking and slow-spiking units based on their trough-to-peak times, which has been shown to indicate the presence of inhibitory and excitatory neurons respectively [39].

The FBR in brain tissue implanted with MEAs is typically quantified using endpoint immunohistochemical (IHC) characterization. The biomarkers used often include glial fibrillary acid protein (GFAP) to mark the development of activated astrocytic glial cells and neuronal nuclei antigen (NeuN) to identify healthy neurons. Immune responses in brain tissue have been found to be directly proportional to the cross sectional surface area of the penetrating shanks of the implanted MEAs [40]–[42]. We hypothesized that the use of ultrathin a-SiC microelectrode arrays will minimize the FBR in a chronic study and will be associated with a relative stability in the recording capability of extracellular action potentials. In this study, we evaluated the electrochemical properties and extracellular action potential recording performance of a-SiC devices implanted in rat motor cortex over 16 weeks. We also measured the mean fluorescence intensity and cell densities derived from GFAP and NeuN images, respectively, from rat brain tissues implanted with a-SiC MEAs and compared these with cortical tissue samples from sham surgeries.

## Materials and Methods

### Microelectrodes

Amorphous silicon carbide (a-SiC) probes were fabricated at The University of Texas at Dallas using standard semiconductor fabrication techniques. For the first step of fabrication, prime grade 100 mm diameter, 525 µm thick Si wafers (Silicon Valley Microelectronics, Inc., USA) were coated with a 1 µm layer of polyimide PI2610 (HD Microsystems, USA). This layer serves as a release layer that, when hydrated in deionized water after fabrication, releases the completed devices from the Si wafer. The PI was cured at 350 °C for one hour under nitrogen gas at atmospheric pressure. Next, 6 µm of a-SiC was deposited using plasma enhanced chemical vapor deposition (PECVD). The PECVD deposition conditions were substrate temperature of 350 °C, pressure of 1000 mtorr, and 270 W RF power, with 164 sccm Ar, 36 sccm CH_4_, and 600 sccm 2% SiH_4_ serving as the reactive gasses in the plasma. A trilayer Ti/Au/Ti (30/150/30 nm) metal interconnect layer from the bond pad region and the electrode sites was patterned using lift-off photolithography following deposition by electron beam evaporation (CHA Industries, USA). Once the metallization layer was completed, a final 2 µm layer of a-SiC was deposited over the metallization and first layer of a-SiC to complete the stack. Next, 2×5 µm^2^ vias were opened in the top a-SiC layer by inductively coiled plasma (ICP) etching (PlasmaTherm, USA) to expose the underlying metallization at the electrode sites. ICP etching was performed with 25 sccm SF_6_ and 5 sccm O_2_ as reactive gasses. In the same step, 200×100 µm^2^ openings for the bond pad region were created at the distal bond pad sites. Sputtered iridium oxide film (SIROF) was used on the electrode sites to reduce electrode impedance and increase charge injection capacity. SIROF was deposited using DC magnetron sputtering as described by Maeng et al. [43]. The SIROF electrodes were circular with a geometric surface area (GSA) of 200 µm^2^ and thickness of 250 nm. A 30 nm sputtered Ti adhesion layer was deposited prior to the SIROF. Finally, fabrication of devices was completed by singulation of the outline of the devices from the stack using ICP etching as described above. Once thin film fabrication was complete, the wafers were soaked in 87 °C DI water for approximately 24 hours to hydrate the base polyimide layer to release completed devices from the silicon wafer. The devices were bonded to connectors (Omnetics Connector Corporation, USA) and 30G stainless-steel reference and ground wires were both attached via H20E conductive silver epoxy (Epoxy Technology, Inc., USA). Lastly, the bond pad area was coated in a medical epoxy (EA M-121HP, Henkel Loctite, Germany) to encapsulate the bond pads and connector. Cross sectional diagrams of the device are shown in figure 1A. Optical and scanning electron microscope images of the completed device and a recording electrode site are shown in figures 1B-E.

**Figure 1:**
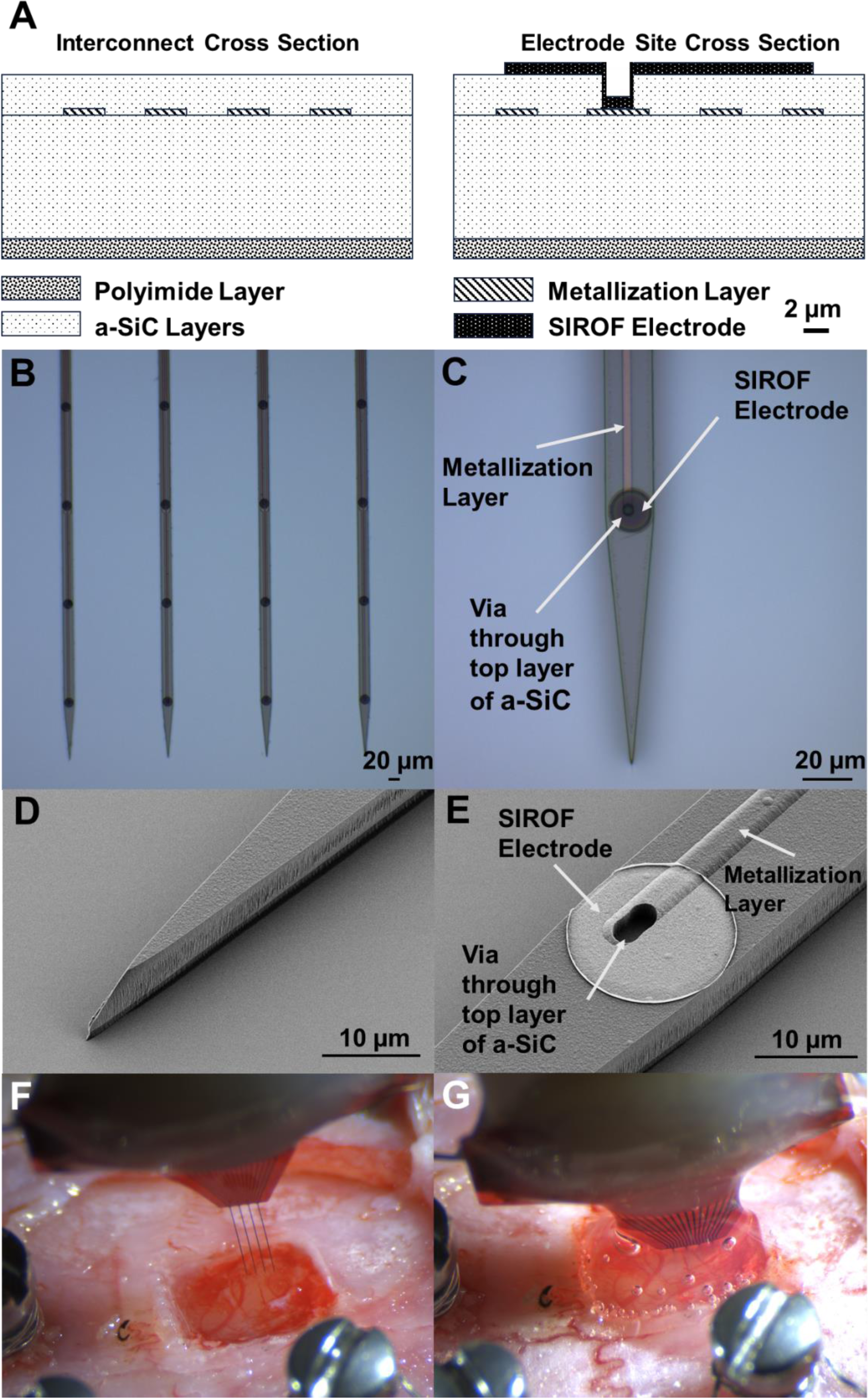
(A) Cross sectional diagram of an a-SiC device. The metal traces that lead along the shank are pictured on the left diagram and the recording electrode site is shown in the right diagram. Diagram is drawn to scale. (B) Optical image of four a-SiC penetrating shanks and (C) detailed image of a single electrode site. The interconnect metallization can be seen in B. SEM of the penetrating tip (D) and a recording electrode site (E). The SIROF film (bright disk) is connected to the underlying metal trace by a via etched through the top layer of silicon carbide. Note that there is an observable 1 µm thick polyimide layer below the a-SiC stack. Before (F) and after (G) implantation of an a-SiC electrode array into rat motor cortex. The size of the craniotomy shown is approximately 2×2 mm. In G, a topical adhesive has been added to close the craniotomy.

These devices had four 2 mm long colinear shanks with four 200 µm^2^ recording sites per shank, for a total of 16 channels per device. Each shank was pitched 200 µm apart from each other and electrode sites were located at 100, 300, 500, and 700 µm from the distal tip of the array. Each shank was patterned with a penetrating tip with an included angle of 11 degrees.

### Surgical Implantation

All procedures were approved by The University of Texas at Dallas Institutional Animal Care and Use Committee (protocol #18-13). Adult female Sprague Dawley rats were used in the study. Initial anesthesia was induced using 2-3% isoflurane in oxygen followed by an intraperitoneal injection of a ketamine (65 mg/ml), xylazine (13.3 mg/ml), acepromazine (1.5 mg/ml) cocktail. A surgical plane of anesthesia was confirmed with tail and toe pinches. The animals were then transferred to a stereotactic frame and anesthesia was maintained with 2% isoflurane in oxygen. A warming pad (Kent Scientific Corporation, Torrington, CT, USA) was placed under the rat and kept at 37 °C. A midline incision was made on the scalp and tissue resected to expose the skull. Stainless-steel bone screws were drilled (Stoeltin Co., Wood Dale, IL, USA) adjacent to the site of implantation, and a ∼2×2 mm craniotomy and durotomy was performed over the left motor cortex (M1) (2 mm anterior from bregma and 2 mm lateral from the midline). Ground and reference wires were wrapped around the bone screws, and devices were implanted with a NeuralGlider Cortical Neural Implant Inserter (Actuated Medical, Inc., USA), which supplied axial vibration in the direction of insertion to aid penetration into the brain. Arrays were inserted at a speed of 100 µm/s to a depth of ∼1.5 mm with vibrational power of 0.5 W. Fig 1F shows the a-SiC device prior to implantation through the craniotomy whereas Fig 1G depicts the same array immediately following successful insertion. After implantation, a collagen-based dural graft was used to replace the resected dura, and silicone elastomer adhesive (Kwik-Sil, World Precision Instruments, Sarasota, FL, USA) was applied to cover the craniotomy. Finally, dental cement was used to form a headcap that encapsulated the device, bone screws, and exposed skull, while leaving the connector accessible. Implanted animals were individually housed and maintained for 16 weeks on a 12-hour light-dark schedule.

### Electrochemical Evaluation

We quantified the electrochemical stability of the implanted electrodes using electrochemical impedance spectroscopy (EIS) and cyclic voltammetry (CV) prior to implantation and throughout the 16 weeks of implantation. EIS measurements were made over a 1-10^5^ Hz frequency range using a 10 mV RMS sinusoidal voltage signal about the equilibrium potential of the electrode. Measurements were made at 10 data points per decade of frequency vs a Ag|AgCl reference electrode. In this study, we report the impedance magnitude at three frequencies, 1 Hz, 1 kHz, and 30 kHz. These frequencies were chosen to identify EIS regions of interest associated with local field potential recording, single unit action potential recording, and the tissue impedance, respectively [44], [45]. CV measurements involved sweeping the electrode potential from +0.8 V to −0.6 V versus Ag|AgCl at sweep rates of 50 mV/s and 50,000 mV/s to monitor electrode functionality and any faradaic reactions occurring at the electrode site [45], [46]. Pushing electrode potentials beyond this potential range may result in irreversible oxidation or reduction of water, which may damage the electrode site or adjacent tissue [46]. The cathodal current over a single CV cycle was integrated over time to give the cathodal charge storage capacity (CSC_c_). Both saline and *in vivo* EIS and CVs were performed using a Gamry Reference 600 potentiostat (Gamry Instruments Inc, Warminster, PA, USA). All electrochemical measurements were performed in a 3-electrode configuration with each electrode site as the working electrode, a large Pt wire counter electrode, and Ag|AgCl reference electrode.

Prior to implantation, CV and EIS measurements of each electrode were obtained in an inorganic model of interstitial fluid (mISF) saturated with CO_2_/O_2_/N_2_ mixed gas (5/6/89% respectively). The mISF measurements allowed for the verification of the functionality of each electrode channel and provided baseline electrochemical properties of the devices prior to implantation. For *in vivo* CV and EIS measurements, Pt counter and Ag|AgCl reference electrodes were taped to the rat’s tail and covered in a phosphate buffered saline soaked gauze to ensure electrical connection to the animal. *In vivo* electrochemical measurements began at 7-13 days post-implantation, and then weekly up to 16 weeks.

Establishing the availability of functional electrodes was a key component in determining active electrode yield, which we defined as the percentage of available electrode sites showing at least one discernable single unit. Electrode site connectivity was assessed from the CSC_c_ determined at slow and fast sweep rates. If the CSC_c_ for the 50 mV/s sweep CV for a given channel was below 1 mC/cm^2^ and the 50,000 mV/s sweep CSC_c_ was below 0.1 mC/cm^2^ the channel was presumed disconnected. The CVs with these low CSC_c_ values were characterized by a lack of observable oxidation-reduction peaks associated with the SIROF. These channels were included in the data analysis for the weeks in which a characteristic *in vivo* SIROF CV was observed. In some cases, a channel was observed to be “disconnected” for one week and connected for subsequent weeks. In this circumstance, the data for the week of observed electrochemical disconnects were not included for electrochemistry analysis, but these channels were considered connected when calculating active electrode yield for neural recording.

### Neural Recordings

On the day of surgery, and each week thereafter with the rats anesthetized with 2-3% isoflurane, wideband (0.1-7000 Hz) neural recordings were collected for 600 s at a 40 kHz sampling frequency using a Plexon acquisition system (Omniplex, Plexon, Inc., USA). To discriminate single units, data were bandpass filtered using a 300 Hz to 3000 Hz Butterworth filter. Common median referencing was used to reduce the level of noise and the effect of artifacts [47]. Next, a −4σ (standard deviation) threshold of the filtered signal was used to extract waveforms that could be extracellular action potentials. Single units were then automatically identified using a k-means scan and manually verified based on separation in principal component space. Signal-to-noise ratio (SNR) was calculated by dividing the peak-to-peak voltage (Vpp) of the discriminated single units by the noise. The noise value was determined by removing any portion of the filtered continuous signal that was associated with a single unit, then calculated as the RMS of the remaining values. Active electrode yield (AEY) for a recording session was calculated as the percentage of available channels that recorded at least one single unit in that session. The spike rate was calculated as the inverse of the median time between spikes associated with a given single unit.

Finally, the trough-to-peak time of each identified single unit was measured. Then, units were further classified as fast-spiking if the spike width was less than 0.4 ms, and as slow-spiking otherwise, based on previously reported literature [39]. This threshold was further validated in this study by analyzing the histogram distribution of spike widths and using the *otsuthresh* function included in MATLAB, which determines a threshold that can be used to separate a bimodal histogram into two groups (Mathworks, Natick, MA, USA). The proportion of slow spiking units with respect to fast spiking units was calculated as the percentage of all single units with a trough-to-peak time greater than 0.4 ms. Electrode channels that were identified as non-functioning based on electrochemistry as described above were excluded from the analysis of recording performance. Single units based on less than 100 waveforms in a 10 minute recording were also excluded from analysis.

### Immunohistochemistry

Immunohistochemical analysis of rat brain tissues implanted with the a-SiC MEAs was performed as described by our group previously [48], [49]. Briefly, following 16 weeks of neural recordings and electrochemical measurements, rats were euthanized by intraperitoneal injection of sodium pentobarbital (Virbac Corporation, Westlake, TX, USA) and transcardially perfused with 350 mL of 1x PBS followed by 300 mL of 4% paraformaldehyde (PFA) (Sigma-Aldrich, Saint Louis, MO, USA). Probes were removed following perfusion. The rat brains were then extracted and submerged in 4% PFA for 48 hours. Then, the brains were marked with a needle approximately 3 mm from the presumptive implant site. Tissue sections were then trimmed to a ∼ 6 mm x 6 mm region centered about the implant site and the tissue block was embedded in a 4% agarose gel (Sigma-Aldrich, Saint Louis, MO, USA). This tissue-gel block was glued onto the slicing platform of a vibratome (VT 1000S, Leica vibratome, Wetzlar, Germany) and sliced in 100 µm thick sections. Slices were then maintained in PBS with 0.1% (w/v) sodium azide (Sigma-Aldrich, St. Louis, MO, USA) at 4°C until staining was performed. Slices were grouped by depth from the cortical surface: superficial (0-400 µm), middle (500-800 µm), and deep (900-1200 µm).

For staining, slices were blocked with 4% (v/v) normal goat serum (Abcam Inc., Cambridge, UK) and permeabilized using 0.3% (v/v) Triton X-100 in 1x PBS with 0.1% sodium azide (Sigma-Aldrich, St. Louis, MO, USA) for an hour. The slices were then incubated overnight at 4°C with primary antibody solutions (supplemented with 3% (v/v) Triton X-100 in 1× PBS) to target astrocytes (GFAP) and neuronal nuclei (NeuN) (Abcam Inc., Cambridge, UK) (Table 1). GFAP staining relied on a 1:500 dilution of the primary antibody. The secondary antibody, goat anti-chicken IgY (Alexa Fluor 647), was prepared at a 1:4000 dilution. NeuN staining also relied on a 1:500 dilution of the primary antibody and a 1:4000 dilution of the secondary antibody, in this case goat anti-rabbit IgG (Alexa Flour 555) (Abcam Inc, USA). After 24 hours, the brain slices were washed thoroughly with 1x PBS, treated with Image-iT fixation/permeabilization kit (Thermo Fisher Scientific), and incubated with a blocking buffer of normal goat serum containing secondary antibodies for 4 hours. The slices were then mounted onto glass slides with Fluoromount aqueous mounting medium (Sigma-Aldrich, St. Louis, MO, USA). Stained slices were imaged using confocal microscope (Model AIR, Nikon Instruments, USA). A maximum intensity projection image was obtained with 5µm Z-steps over a total range of 35 µm using Nikon NIS elements software (Nikon Instruments, USA). Hardware/software settings were conserved across image acquisition. The objective was set to 10x, and images were obtained over a 5000 x 5000 µm viewing window.

The GFAP intensity of an image was calculated using custom MATLAB code. Briefly, 100 x 50 µm regions-of-interest (ROIs) were created on opposite sides of holes identified as probe locations, with the 100 μm length of the ROI oriented in parallel with respect to the shank orientation. The intensity within these rectangular regions was then calculated. Each of the regions were then shifted 50 µm away from the previous location, perpendicular to the shank orientation. This extended up to 500 µm away from the holes and was repeated for each hole location. Data were normalized to the intensity measured in the rectangular region at 450-500 µm away from each hole. Next, the GFAP intensity of all the slices for a given distance from the marked implant site within the superficial region (100-800 µm from the surface) and deep region (800-1200 µm from the surface) were averaged for further statistical analysis. To identify neuronal cell bodies in proximity to the implant site, cells with a response to NeuN staining were counted manually in similar 100 x 50 µm rectangular regions. Cell totals for each region were summed up and again normalized to the region at 450-500 µm away from each hole. In total, seven a-SiC implanted brains were processed, with the superficial region consisting of 18 slices and the deep region consisting of 8 slices. There are instances where implantation holes could not be readily identified for a-SiC probes, especially at depth. These slices were included in the analysis, but the hole location was estimated through corresponding slices from superficial layers. In total, 5 sham brains were processed, consisting of 13 slices for the superficial and 10 slices for the deep layers.

Following the acquisition of brain slice images, we observed an edge effect in fluorescence intensity distribution, with higher intensity in the center of the images. This variation in fluorescence intensity was determined to be primarily due to an imaging artifact rather than a true representation of higher fluorescence intensity in the central regions. We confirmed this observation in two ways: First, the edge effect appeared across all the acquired images. Second, moving the microscope stage and centering the regions near the previously scanned region on the same slice showed the same edge effect. We hypothesize that this edge effect is due to tissue slice’s thickness. To address this issue, we performed a gradient adjustment by calculating the linear regression for each of the sham images and subtracted it from the original data. Then, we applied the same subtraction to the contralateral images with a-SiC implants to adjust for the observed gradient. While the edge effect was observed on both NeuN, and GFAP images, this adjustment was only applied to GFAP images as its analysis was based on intensity, while NeuN analysis was based on single cell counting, which was performed without affecting the analysis.

### Statistical Analysis

Normality of data sets was tested with the Shapiro-Wilk’s normality test and verified using QQ plots. All electrochemical data sets were non-normal (Shapiro-Wilk, *p* < 0.05), with a skewness toward larger values for both impedance and CSC_c_ values. Consequently, electrochemical data are reported as: median, quartile, and min-max range. To analyze differences in mISF and *in vivo* impedance and CSC_c_ following implantation, we performed a Mann-Whitney *U* Test. To assess electrode characteristics over the time-course of implantation, we performed a robust linear regression on the *in vivo* time series data (Stata 18, College Station, TX, USA) and reported the deviation from zero of the resulting slopes of the linear fit. Neural recording data were also non-parametric (Shapiro-Wilk, *p* < 0.05). Recording data are also presented as median, quartile and range with changes over the 16-week study period assessed by a robust linear regression. Day of surgery recording data were excluded from the linear regression due to possible noise artifacts arising from the surgical suite as well as a difference in anesthesia between implant surgery and subsequent recording sessions. Immunohistochemical data were normally distributed (Shapiro-Wilk, *p* > 0.05). We report the mean and standard error for each of the IHC measurements. To understand the effect of distance from the implantation site on the two IHC metrics, we performed a paired *t*-test between the metric of interest at one binned distance of a-SiC implanted slices and the same distance of the sham slices.

## Results

### Electrochemistry

EIS measurements of implanted MEAs demonstrated marked consistency over the chronic implantation period. A representative example of the changes in impedance modulus from one electrode site prior to implantation (mISF) and then at 1 week, 6 weeks, 12 weeks, and 16 weeks post-implantation is shown in Figure 2A. For all animals, trendlines of median, quartiles, and ranges for impedance magnitudes at frequencies of 1 Hz, 1 kHz, and 30 kHz are shown in Figures 2B, 2C, and 2D, respectively. Pre-implantation median mISF impedance is plotted and shown as a grey box and whisker plot. Median impedance in mISF was 10.8 MΩ (7.2 MΩ, 19.1 MΩ) (1^st^ quartile, 3^rd^ quartile) at 1 Hz, 0.23 MΩ (0.14 MΩ, 0.42 MΩ) at 1 kHz, and 0.049 MΩ (0.032 MΩ, 0.104 MΩ) at 30 kHz. Post-implantation, the median 1 Hz impedance of 11.6 MΩ (8.2 MΩ, 20.2 MΩ) was essentially unchanged from the pre-implantation level in mISF and did not show a significant difference between mISF and cortex (*p* = 0.556, Mann-Whitney *U* Test). Median 1 kHz and 30 kHz impedance showed a significant, and not unexpected, increase in impedance between mISF and week 1 of implantation. The week 1 median impedance at 1 kHz was 1.2 MΩ (0.71 MΩ, 1.6 MΩ) and was significantly different (*p* < .0001, Mann-Whitney *U* Test) from mISF. Likewise, the week 1 median 30 kHz impedance was 0.11 MΩ (0.10 MΩ, 0.13 MΩ) and was significantly different from mISF (*p* < .0001, Mann-Whitney *U* Test). *In vivo*, median 1 Hz impedance increased moderately over 16 weeks to 13.5 MΩ (9.99 MΩ, 46.1 MΩ). Median impedance at 1 kHz increased to 2.9 MΩ (0.74 MΩ, 3.55 MΩ) and median 30 kHz impedance increased to 0.13 MΩ (0.12 MΩ, 0.15 MΩ) at week 16. Overall, the observed changes in impedance post-implantation were modest, although, as shown in figure 2A, there was variability, particularly in the mid-frequency range (10^2^-10^4^ Hz). A robust linear regression analysis did not reveal a time-dependent change in the 1 Hz impedance over 16 weeks (*p* = 0.421, slope = −0.034 MΩ/week, 95% CI = (−0.11 MΩ/week, 0.049 MΩ/week)). For the 1 kHz data, we observed a small but significant positive trend (*p* < 0.001, slope = 0.059 MΩ/week, 95% CI = (0.043 MΩ/week, 0.073 MΩ/week)); however, it is not expected that the magnitude of the change would compromise the recording performance of the electrodes. Likewise, we observed a small but significant positive trend in the 30 kHz impedance (*p* < 0.001, slope = 0.001 MΩ/week, 95% CI = (0.0008 MΩ/week, 0.0015 MΩ/week)). While the 1 kHz and 30 kHz trendlines are significant, the small magnitude of the slope of the best fit line is indicative of the stability of these probes over time while *in vivo*.

**Figure 2:**
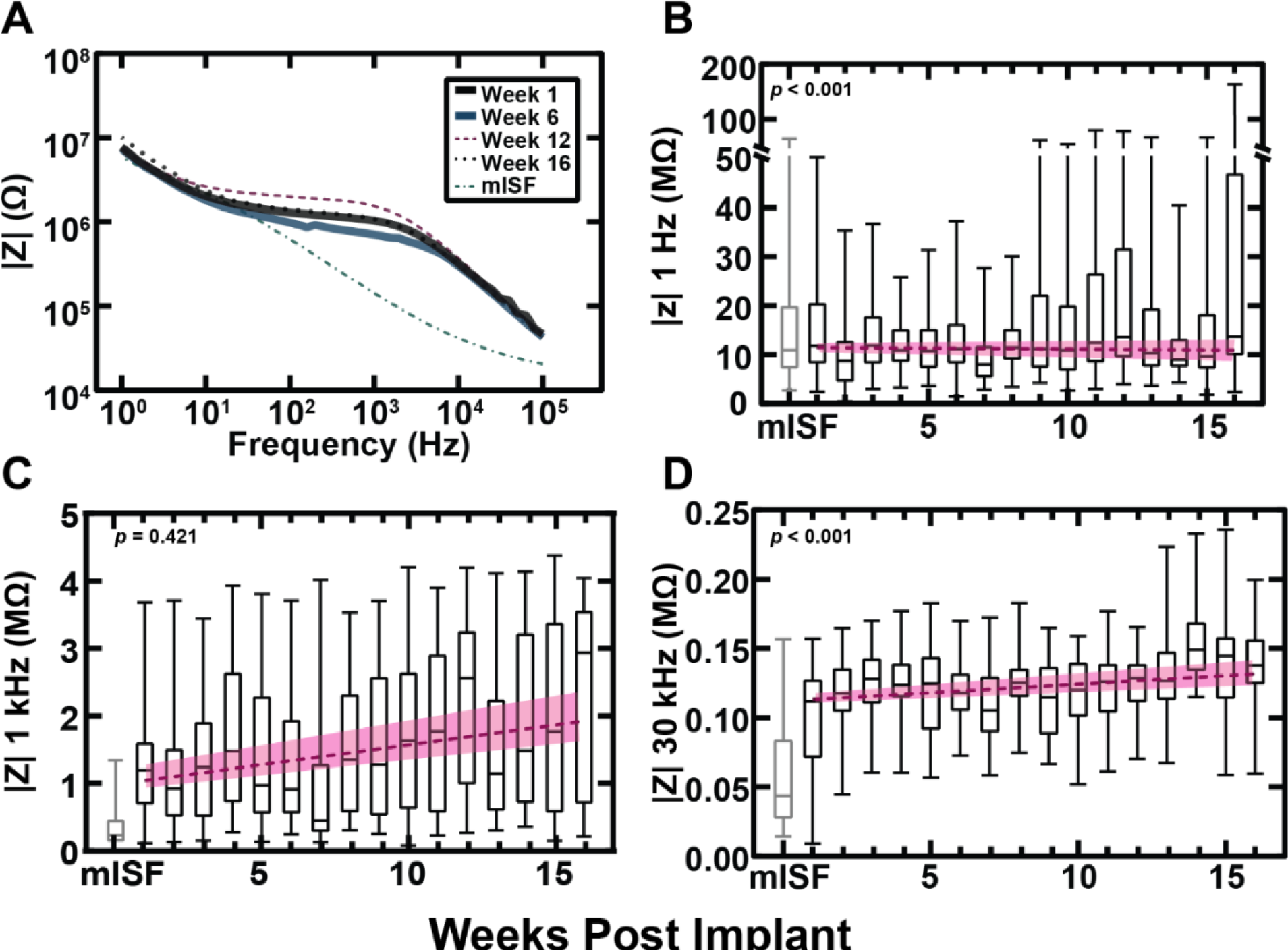
(A) Representative impedance modulus traces from a single SIROF electrode site on a single array in mISF and at 1-, 6-, 12-, and 16-weeks post implantation. (B, C, D) Median, quartiles, and range for impedance magnitude trends at frequencies of 1 Hz (B), 1 kHz (C), and 30 kHz (D). Each point represents the median of all available electrodes across all arrays for every week. The grey box and whisker plot at week 0 represents the results from *in vitro* mISF measurements. Note the scale break in the |Z| axis for B. A robust linear regression was fitted to each of the data sets. *p* values for the slope’s deviation from 0, calculated slope, and 95% CI are as follows: 1 Hz impedance: *p* = 0.421, slope = −0.034 MΩ/week, 95% CI = (−0.11 MΩ/week, 0.049 MΩ/week). 1 kHz impedance: *p* < 0.001, slope = 0.059 MΩ/week, 95% CI = (0.043 MΩ/week, 0.073 MΩ/week. 30 kHz impedance: *p* < 0.001, slope = 0.001 MΩ/week, 95% CI = (0.0008 MΩ/week, 0.0015 MΩ/week).

CV measurements also revealed chronic stability similar to the impedance data. Representative CV curves from a single SIROF electrode at 50 mV/s and 50,000 mV/s sweep rates are shown in figures 3A and 3B, respectively, noting the difference in current scales between 50 mV/s and 50,000 mV/s. Figures 3C and 3D shows the median CSC_c_ for each sweep rate over 16 weeks, including a robust regression fit to the data. Median and quartile CSC_c_ values in mISF were 53.2 mC/cm^2^ (37.3 mC/cm^2^, 71.5 mC/cm^2^) at 50 mV/s and 7.22 mC/cm^2^ (4.12 mC/cm^2^, 11.1 mC/cm^2^) at 50,000 mV/s. On the first week following implantation, the median CSC_c_ at 50 mV/s increased to 58.1 mC/cm^2^ (35.8 mC/cm^2^, 76.2 mC/cm^2^). At 16 weeks post-implantation, the median 50 mV/s CSC_c_ was 55.9 mC/cm^2^ (49.1 mC/cm^2^, 64.2 mC/cm^2^). The mISF and week 1 50 mV/s CSC_c_ were not significantly different (*p* = 0.574, Mann-Whitney *U* Test). Like the impedance data, we fit a robust linear regression model to the *in vivo* reported CSC_c_ data. A robust linear regression showed no significant trend in 50 mV/s CSC_c_ over 16 weeks ((*p* = 0.112, slope = 0.237 mC/cm^2^-week, 95% CI = (−0.055 mC/cm^2^-week, 0.528 mC/cm^2^-week). At one-week post-implantation, the 50,000 mV/s CSC_c_ declined to 4.27 mC/cm^2^ (2.97 mC/cm^2^, 6.61 mC/cm^2^), which was a significant difference (*p* < .0001, Mann-Whitney *U* Test). Subsequently, the median 50,000 mV/s CSC_c_ showed a slight but significant increase reaching 5.93 mC/cm^2^ (4.01 mC/cm^2^, 7.72 mC/cm^2^) by week 16. We observed a significant positive slope in the faster 50,000 mV/s sweep CSC_c_ (*p* < 0.001, slope = 0.063 mC/cm^2^-week, 95% CI = (0.028 mC/cm^2^-week, 0.098 mC/cm^2^-week). These changes in CSC_c_ over 16 weeks, are extremely small and suggest that the SIROF electrode coatings and encapsulation are generally stable *in vivo* over this implantation period. Overall, these electrochemical observations suggest relatively consistent stability for chronically implanted a-SiC devices.

**Figure 3:**
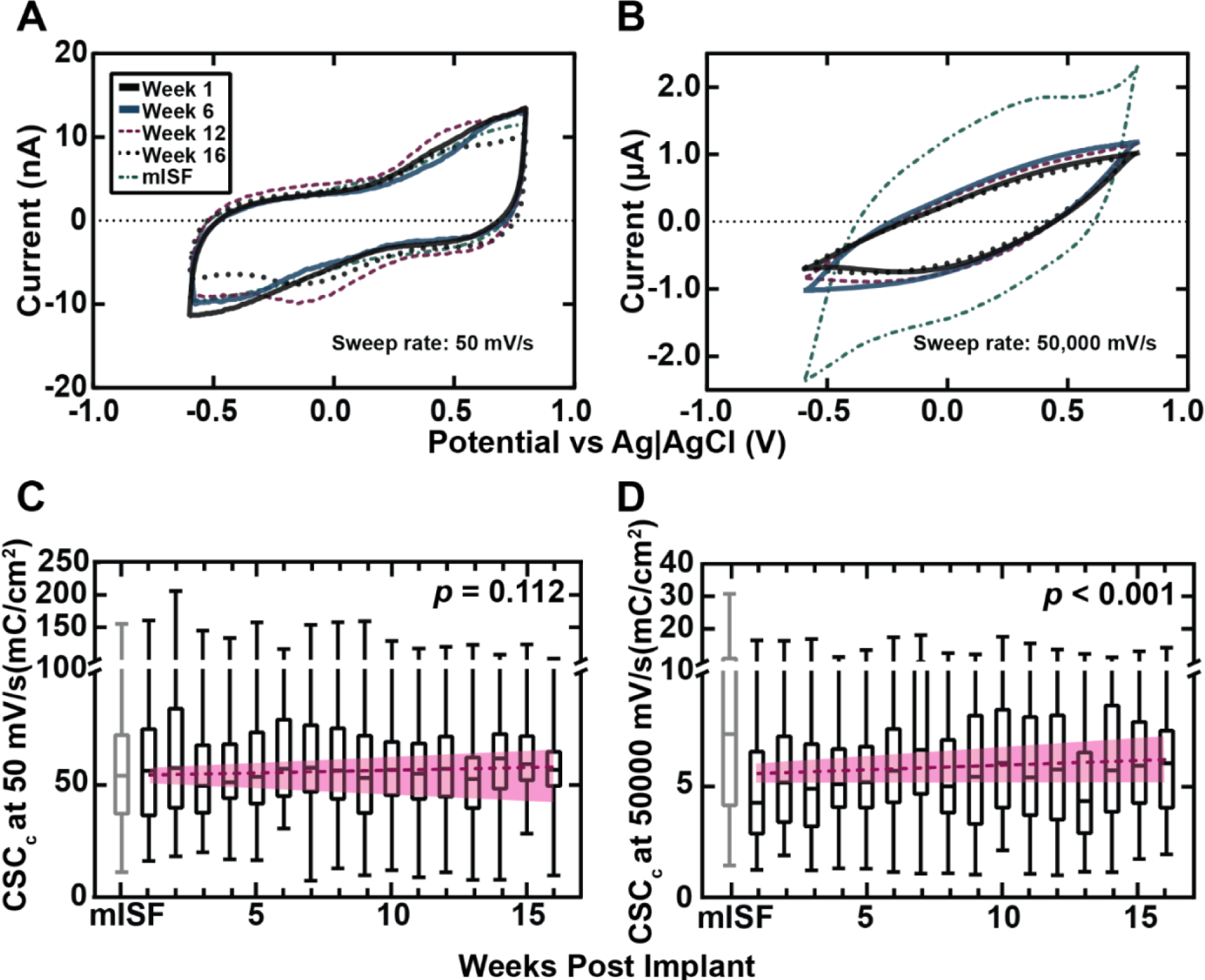
Representative CV curves at 50 mV/s (A) and 50,000 mV/s (B) sweep rate of a SIROF electrode site in mISF and at weeks 1, 6, 12, and 16 post implantation. (C, D) Median, quartiles, and range for CSC_c_ at 50 mV/s (C) and 50,000 mV/s (D). Box and whisker plots show the median CSC_c_ of all available electrodes across all arrays for that week. Note the scale break in the CSC_c_ axis for C and D. A robust linear regression is fitted to the data and plotted with a 95% confidence interval. Pre-implantation mISF data were omitted from the regression analysis. The grey box and whisker plot at week 0 is from mISF measurements. *p* values for the slope’s deviation from 0, calculated slope, and 95% CI are as follows: 50 mV/s sweep rate: *p* = 0.112, slope = 0.237 mC/cm^2^-week, 95% CI = (−0.055 mC/cm^2^-week, 0.528 mC/cm^2^-week. 50,000 mV/s sweep rate: *p* < 0.001, slope = 0.063 mC/cm^2^-week, 95% CI = (0.028 mC/cm^2^-week, 0.098 mC/cm^2^-week).

### Neural Recordings

A-SiC devices recorded neural activity over the 16 weeks of implantation within rat motor cortex. Figure 4C shows the AEY from all a-SiC MEAs. Each point represents the collection of all recordings taken each week post-implantation, across all functional electrode channels in all animals. On the day of surgery for each animal, all 112 electrode channels appeared functional, and 92% recorded single unit activity. 16-weeks post-implantation, the AEY declined to ∼51% of the 80 remaining functioning channels, indicating that single units could still be detected on a small majority of channels.

**Figure 4:**
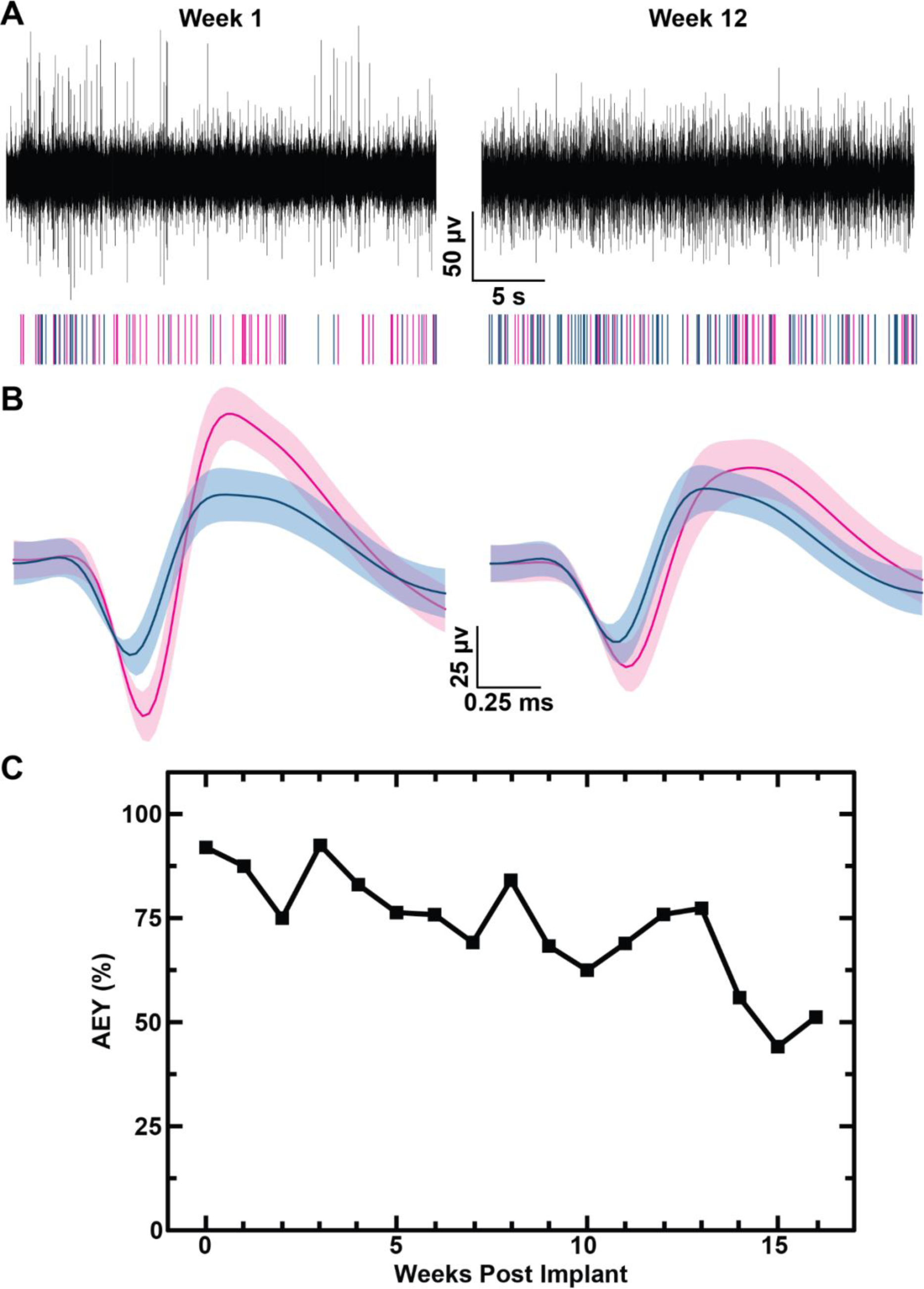
Representative single unit neuronal recordings, unit identification, and AEY over time. (A) representative 30 s epoch of band-pass filtered neural activity (300-3000 Hz) with a corresponding raster plot showing spikes associated with two representative single units in the epoch from weeks 1 (left) and 12 (right) and (B) their corresponding averaged neuronal waveforms, classified as fast spiking (blue) and slow spiking (pink). (C) AEY for each week following implantation.

Figure 5 shows additional electrophysiological data. We observed a small decline in the average peak-to-peak voltage (*p* < 0.05, slope = −1.35 µV/week, 95% CI = (−1.85, −0.85)), beginning at 141 µV on the day of surgery 114 µV 1-week post implantation and a final amplitude of 85 µV at the end of the study. In contrast, we observed stable values in RMS noise as the slope of the trend line was not-statistically significant from zero (*p* = 0.24, slope = 0.022 µV/week, 95% CI = (−0.015, 0.059)). These results yielded a small decline in μV signal-to-noise ratio (−0.19 /week) that was found to be significant (*p* < 0.05, slope = −0.19/week, 95% CI = (−0.24, −0.14)). Finally, the spike rate appeared to have a significant decrease (*p* < 0.05, slope = −0.35 Hz/week, 95% CI = (−0.43, −0.26)).

**Figure 5:**
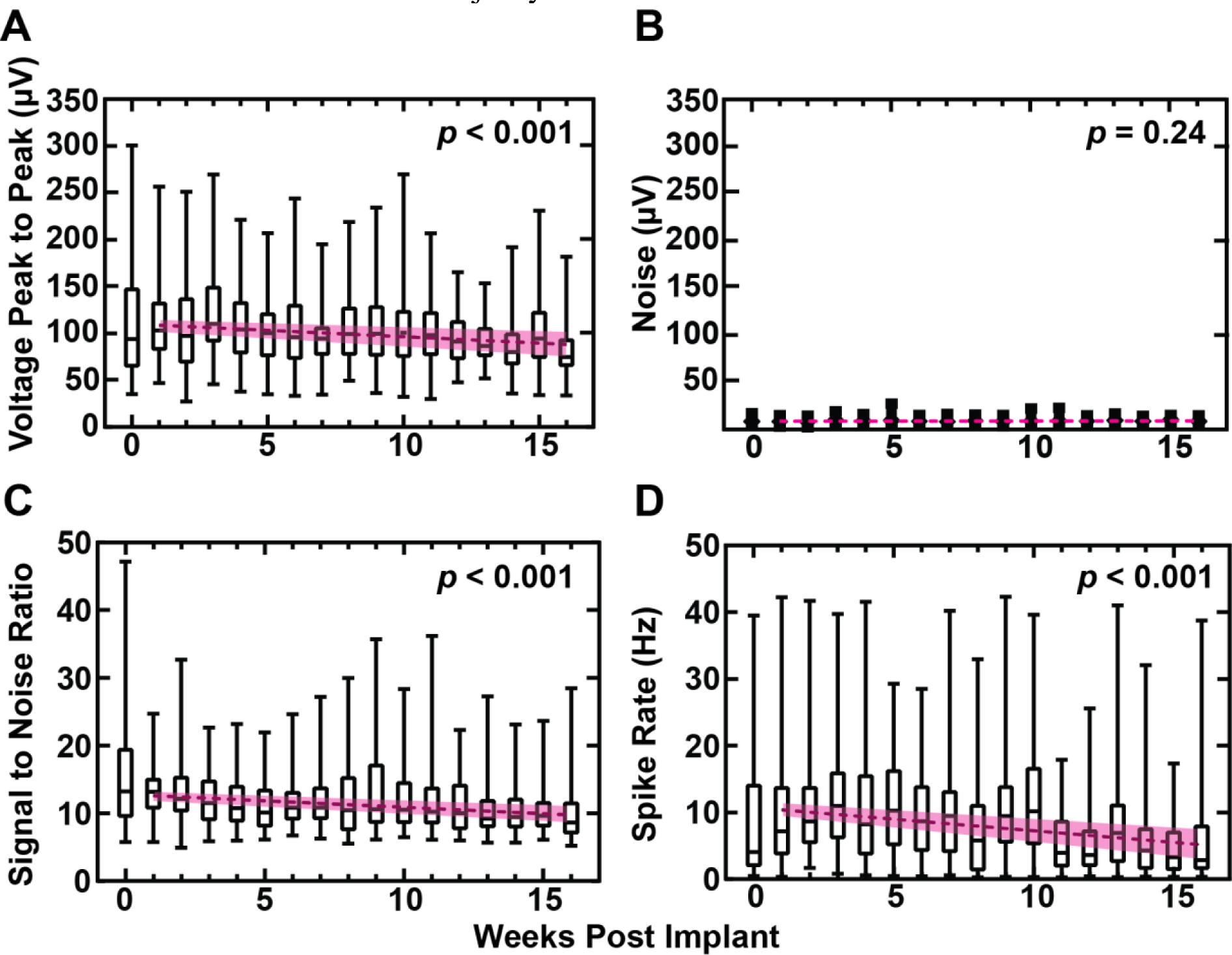
Summary of weekly neural recording data of all functional electrodes across all arrays for day-of-surgery and weeks 1-16: (A) peak-to-peak voltage, (B) RMS noise, (C) signal-to-noise ratio, and (D) spike rate. Boxes represent the 1^st^ and 3^rd^ quartiles, the horizontal line inside each box represents the median, and the whiskers show the range of values from minimum to maximum after removal of outliers. A robust linear regression, omitting day-of-surgery data with 95% confidence intervals is included in pink. Here week 0 represents the neural recording during the day of surgery, but because of the potential effects of surgical anesthesia, it has been excluded from regression analysis.

Figure 6 shows an example of a fast-spiking, or putative inhibitory, neuron and a slow-spiking, or putative excitatory, neuron, defined by having a trough-to-peak time of less or greater than 0.4 ms [39]. The histograms in figure 6B show the number of units in each time bin. The broken line indicates the 0.4 ms threshold at one week and twelve weeks post-implantation. The graph in figure 6C shows the average units per active channel (defined in the section on electrochemistry) that were classified as either excitatory or inhibitory, as well as a robust linear regression line for both putative excitatory (*p* = 0.026, slope = −0.022 units/channel, 95% CI = (−0.041, −0.0030)) and inhibitory (*p* = 0.0024, slope = −0.046 units/channel, 95% CI= (−0.073, −0.019)). Both slopes were significantly different from zero, but they were not statistically different from one another (*p* = 0.13). Figure 6D shows the percentage of neurons we considered slow or fast spiking each week.

**Figure 6:**
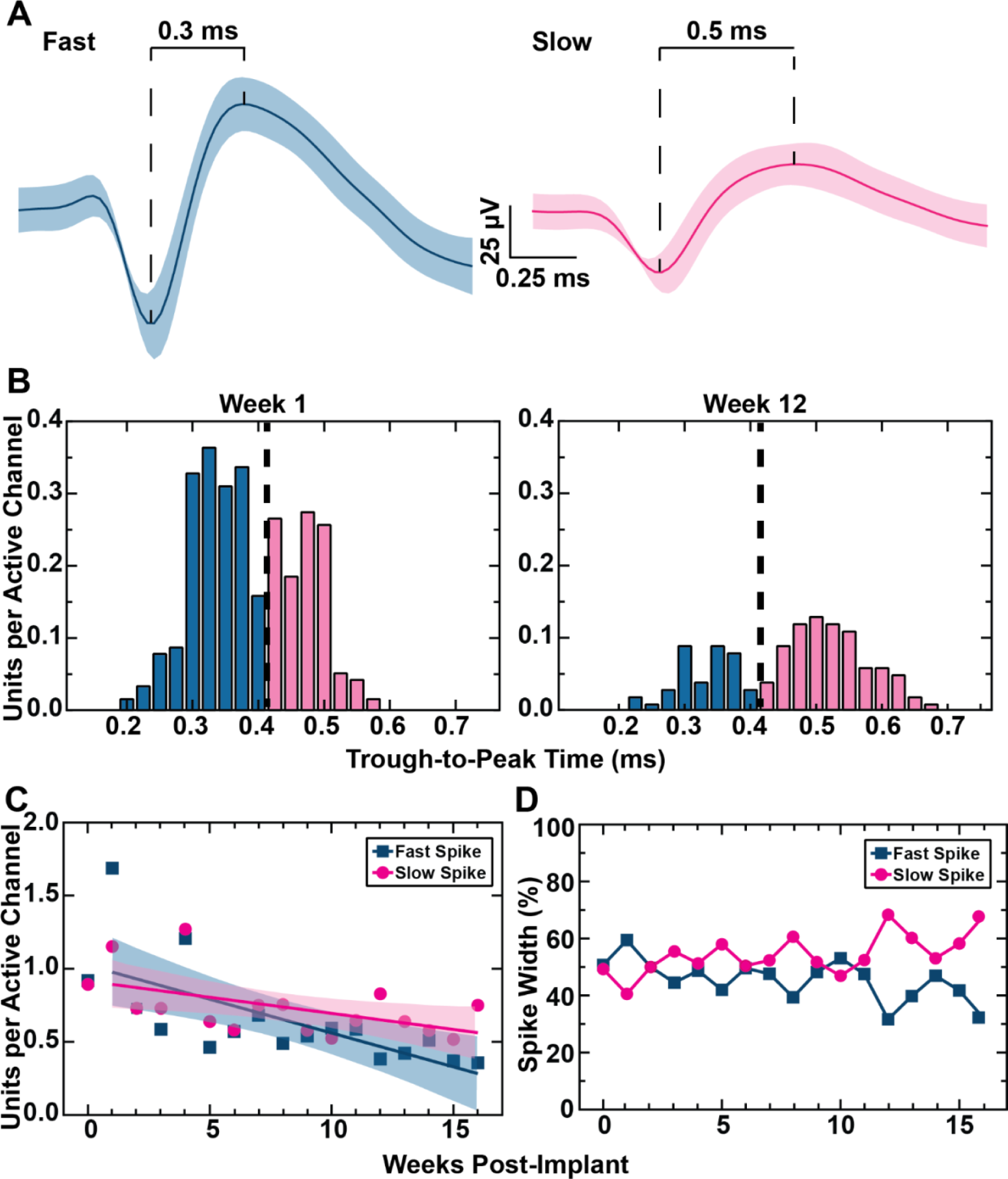
(A) Representative fast (left) and slow (right) spiking units, which are putatively inhibitory and excitatory respectively. (B) Histograms showing the units per active channel classified as fast and slow spiking on week 1 (left) and week 12 (right). The broken line represents the threshold identified using *otsuthresh* as described above. (C) the units per active channel each week classified as fast (blue) and slow (pink) spiking and linear regression for the change of both. There is no statistically significant difference in the rate of decline of either type of unit. (D) the proportion of fast (blue) and slow (pink) spiking units each week.

### Immunohistochemistry

A schema showing the approximate placement of the electrodes with respect to the cortical layers is shown in Figure 7A. Representative stained slices of brains from depths of 100-800 µm and 800-1500 µm for a-SiC devices and contralateral brain slices with no implant (sham) are shown in Figure 7B. These depths were chosen to highlight differences in FBR in the superficial layer, where there are no electrode recording sites, (100 to 800 µm deep) and recording depths where electrode sites reside (800-1500 µm deep). Figure 7C displays the average relative intensity as a function of distance from the implant site (or sham implant site) for GFAP and NeuN at the superficial and electrode site depths. For GFAP staining around the implantation sites, the a-SiC device exhibited an average relative intensity of 1.72 ± 0.29 (mean ± standard error) for superficial depths between 0-50 µm away from the implant site. We observed that the GFAP response decreased to 1.01 ± 0.02 between 450-500 µm away from the implant site. The lack of statistical difference between these two relative intensities suggests no adverse tissue response from the implanted a-SiC array. Also at the superficial depths, sham slices had a mean relative intensity of 1.01 ± 0.02 between 0-50 µm and 1.02 ± 0.02 450-500 µm from the implant site. At 250 µm from the implant site, there was no significant difference between a-SiC slices and sham slices (*p* = 0.326, paired *t*-test). At recording electrode site depths, we observed that the mean relative intensity of a-SiC slices was 1.02 ± 0.02 at 0-50 µm from the implant site, decreasing to 0.93 ± 0.01 at 450-500 µm. For the deep layer (800-1500 µm) sham slices were found to have a mean relative intensity of 0.93 ± 0.02 at 0-50 µm from the implant site and 0.93 ± 0.02 at 450-500 µm. We found that at 150 µm away from the implant site, there was no significant difference between a-SiC slices and sham slices for recording electrode depths (*p* = 0.215, paired *t*-test). These results indicate a depth-dependent astrocytic response that is more prominent at superficial depths, consistent with prior work [49]. The GFAP intensity and spatial extent from the probe site both decreased in the deeper layers.

**Figure 7:**
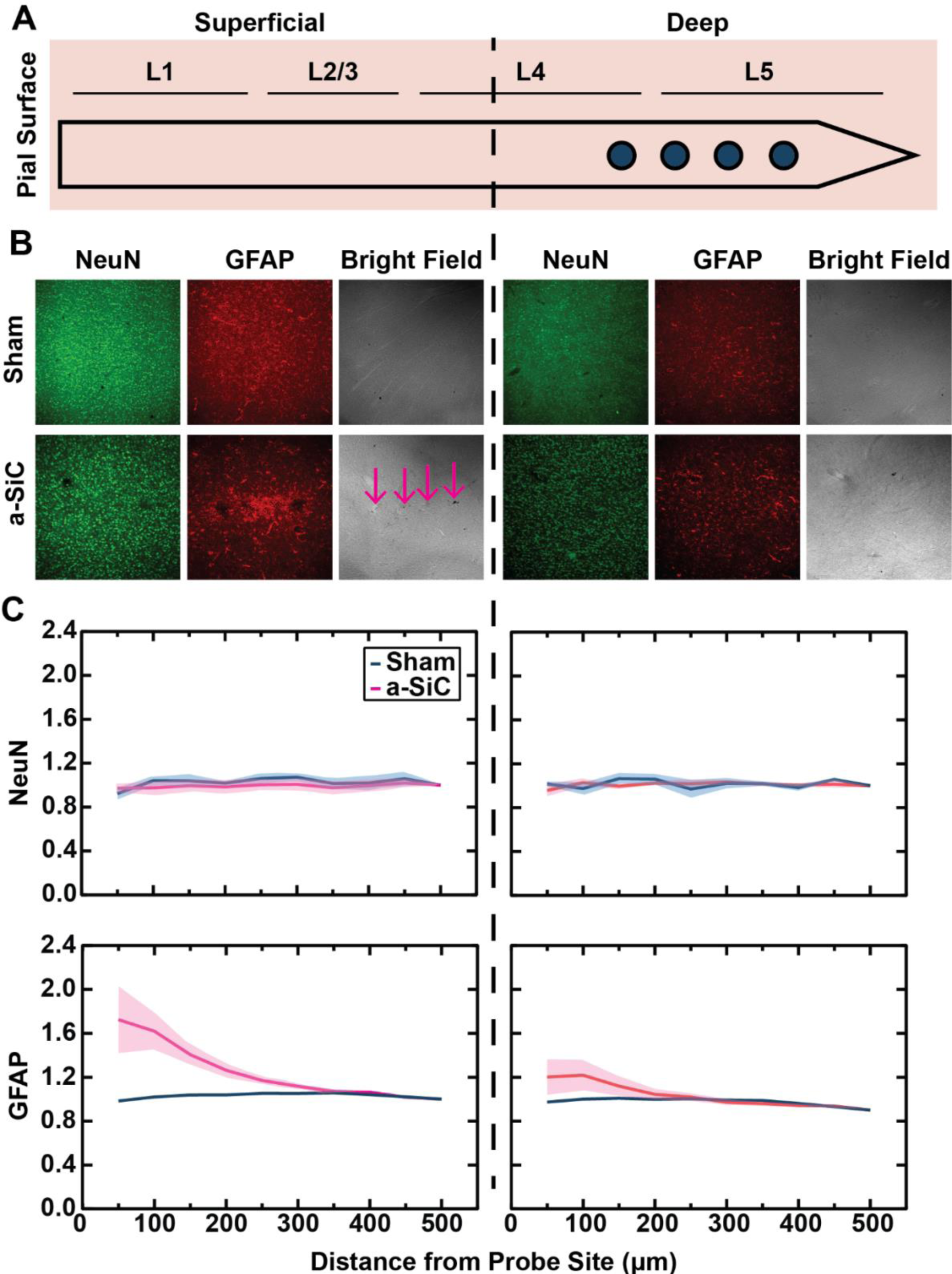
(A) Diagram of one shank of an MEA showing the superficial and recording-depth regions. (B) Representative IHC staining for brains implanted with a-SiC devices from the region surrounding the implant and from the contralateral sham (un-implanted) hemisphere. Images at a superficial region (100-800 µm, left) and deeper (800-1500 µm, right) below the surface of the cortex are shown for GFAP, NeuN, and bright field images. Bright field images were used to identify implant hole locations. (C) Mean and standard error for relative intensities of NeuN and GFAP staining. Normalized NeuN count and GFAP intensity taken from the 100-800 µm depth (left) and from 800-1500 µm depth (right).

As evidenced by analysis of NeuN staining, neural cell counts were not different between implant sites and sham controls for both the superficial and deep depths. At superficial depths, we found that the mean relative intensity of a-SiC implanted slices was 0.92 ± 0.04 at 0-50 µm away from the implant locations and 1.02 ± 0.06 at 450-500 µm. Sham slices were found to have a mean relative intensity of 0.97 ± 0.02 at 0-50 µm and 0.99 ± 0.04 at 450-500 µm. At recording depths, the relative intensity of a-SiC implanted devices was 0.95 ± 0.05 at 0-50 µm and 1.01 ± 0.04 at 450-500 µm. Similarly, sham slices had a mean relative intensity of 1.01 ± 0.02 at 0-50 µm and 1.05 ± 0.01 at 450-500 µm from the pseudo implant location. For both depths, when comparing normalized cell counts at similar distances away from the implant location we found no statistically significant difference between the a-SiC implanted slices and the sham slices (at 0-50 µm, Superficial layers, *p* = 0.352, paired *t*-test; at 0-50 µm, Recording Depth layers, *p* = 0.365, paired *t*-test). From these data we conclude that while a-SiC electrode arrays may cause some astroglial response, they do not cause significant loss of cortical neurons over the 16-week implantation period. Altogether, the results from electrochemistry, neural recordings and immunohistochemistry show stability of the neural interface enabling long-term single neuron recordings with a minimal foreign body response that preserves neurons in the proximity of the implanted MEAs.

## Discussion

In this work, we have demonstrated for the first time that a-SiC devices created using thin film fabrication processes can be used to record neural activity in rat motor cortex up to 16 weeks. In addition, *in vivo* measurements of electrode impedance and cyclic voltammetry were stable over this time period. Given that the linear regression trendlines were not significantly different from zero in slow sweep CSC_c_ measurements, there was little to no evidence of a breakdown of the encapsulation of the insulating a-SiC leading to leakage pathways that might affect the ability to record or possibly stimulate neural activity. The generally flat trend in CSC_c_ for both low (50 mV/s) and high (50,000 mV/s) sweep rates also indicates that there is no consistent access to underlying metallization or significant leakage pathways between channels. This lack of large increases in CSC_c_ for both sweep rates can also be interpreted as a constant shunt capacitance of the dielectric material [50]. A leaky electrode array would have the capacitive dielectric layer become lower impedance over time, increasing the amount of current that can flow through the channel [50]. We do not observe this with the a-SiC devices used in this study, which suggests that a-SiC is a stable dielectric material chronically in animal.

Moreover, the lack of a negative trend suggests that the low impedance SIROF film that was used in these recording electrodes remained intact and stable over 16 weeks in the animal. At the end of the 16-week period, 24% of electrodes were lost due to presumptive fracturing of the metallization trace or to disconnection with the backend connector. Given that both the EIS and CV measurements observed in this study remained relatively consistent over time, a-SiC is a good candidate for electrical insulation and bulk construction of neural interface devices.

Although a-SiC MEAs showed a 40% decline in unit recordings, 51% of connected electrode channels continued to record single unit activity at the end of the study. At week 13, 76% of connected electrode channels were recording single unit activity. Previous observations have reported on the AEY of Si-based electrodes with a decline to less than 10% within just 8 weeks post-implantation [25], [49]. It is important to note that [49] revealed a significant difference in the rate of decline between single- and multi-shank planar Si-based electrodes, where multi-shank devices can experience up to a 90% decline within 8 weeks post-implantation. These observations suggest that the surgical impact of multi-shank implantation and indwelling presence may accelerate the neuroinflammatory response, leading to early failure of the neural interface. This is of importance because a-SiC on planar MEAs reported here showed a slower decline in AEY post-implantation, suggesting that the combination of reduced cross-sectional dimensions and a-SiC insulation can improve the long-term reliability of these devices post-implantation. In addition, our previous study using a-SiC [51] as an insulation material on Utah-style microelectrode arrays observed a comparable decline in the AEY, which further supports the use of these material for intracortical neural interfaces. Other measures of recording capability also showed stability over the course of the study. There was no statistically significant change in noise levels recorded throughout the study. The recording performance of the a-SiC devices was comparable to that of the Si based devices of similar colinear shank design presently used in many neural recording studies [13], [52]. The SNR of recordings made using Si devices has been observed to decline from 8 to 6.8 in a 16-week study [38]. At week 16, recordings made using a-SiC devices had a median SNR of 8.9. We also attempted to distinguish neuron types and examine the relative stability of these neuron types based on the trough-to-peak durations of single unit waveforms. The ability to distinguish neuron types may offer insights into any differential stability of single units as a function of different array geometries or modifications [53]. The number of fast- and slow-spiking neurons per functioning electrode channel declined over time, consistent with the decline in AEY. However, the difference in the rate of decline between fast- and slow-spiking neurons was not statistically significant, indicating both groups are similarly impacted by the implant.

Furthermore, we have demonstrated that a-SiC based probes exhibited minimal FBR, as assessed by reactive astrocyte and neuronal nuclei expression, especially at depths 800 µm below the surface of the pia where the recording electrodes reside. The astroglial scar around a neural implant can begin to form as early as 48 hours post-implantation [21], [54], and the recording performance can diminish with increasing separation between electrodes and presumptive neuronal source [24], [25]. Overall, a-SiC based probes exhibited reduced FBR as compared to Si based arrays when examined qualitatively with a previously reported study, which showed significant astroglial response and decline in local neural cell bodies near the implant site at depths 1400 µm below the pial surface [24]. We have also found that the FBR is similar to that of ultra flexible devices, such as the StimNET, which demonstrated a 10% increase in GFAP expression at 50 µm away from the implanted array versus a control and no significant difference at 100 µm away from the implant location [55]. In this work, we observed a 20% increase over a sham control and no significant difference in GFAP expression at 150 µm away from the implant location. NeuN expression was similar in both studies, with no significant difference between implanted and control slices [55]. Additionally, a-SiC probes in this work had a calculated stiffness of 0.8 mN/m. When compared to the stiffness of devices examined in a meta-analysis in Stiller et al. [56], the stiffness of a-SiC devices are comparable to other devices associated with reduced GFAP expression.

Due to the minimal histological footprint of the a-SiC MEAs, the identification of implant sites in the rat brain tissue specimens was challenging. We analyzed a large, 5000 µm x 5000 µm ROI from the captured histology images around the implant sites of the a-SiC MEAs. However, there were instances where holes generated from implant sites could not be observed, especially from slices below 900 µm from the cortical surface. In these cases, we estimated the location of the a-SiC shanks based on more superficial slices that clearly showed the implant holes. We attribute the absence of an observable FBR in the deep slices to the flexibility of the a-SiC shanks, which likely reduces micromotion between the shanks and brain tissue. The shanks are more flexible at greater depths due to decreased resistance to bending as the implanted shank length increases [32]. Anecdotally, recovered a-SiC probes were found to have minimal tissue adhesion on explanation, when compared to Si-based devices that were at times completely encapsulated in a fibrous tissue [20]. However, a-SiC probes were difficult to consistently retrieve from the cortex to analyze tissue adhesion. Superficial layers of the tissues implanted with a-SiC probes have a higher concentration of activated astrocytes compared to the deeper layers, which, similarly, we associate with a length-dependent reduced flexibility of the shanks close to the pial surface of the brain. Based on the NeuN staining data in Figure 7C, no difference in viability of neurons between the implant and sham sites with either depth or distance from the implant was observed. There was, however, a significantly larger astrocytic response to the implant at superficial layers versus deeper layers. Similar depth-dependent differential expression of GFAP and NeuN was reported by Usoro et al. [49]. While cortical inhomogeneity may play a role in the increased astroglial response in superficial layers [57], there was a reduced response in recording layers where a-SiC devices were their most flexible.

The thin-film fabrication techniques for a-SiC neural interface devices offer several significant benefits. Firstly, the fabrication process for thin-film devices is generally simpler and more accessible compared to devices that must be assembled by hand, for example carbon nanotube devices. The thin film deposition methods used to fabricate a-SiC in this study allow for uniform and precise deposition of a-SiC layers onto a variety of substrates. This, along with the photolithographic patterning techniques, enables the production of a variety of device geometries to meet the needs of the study of interest. Additionally, thin film a-SiC devices are amenable to manipulation and integration with existing electronic systems. The compatibility of a-SiC with traditional silicon-based microfabrication techniques may allow for future active electronic circuitry to be incorporated in the device architecture. In summary, the ease of fabrication and manipulation of thin film a-SiC neural interface devices, along with their reduced FBR activation and stable electrical insulation, makes them a promising neural interface device platform, with the potential for large-scale production and integration into practical brain computer interfaces.

## Conclusion

Minimizing foreign body response and having stable electrode and device materials are two important developments to preserve chronic functionality of neural recording interfaces. In this work, we have demonstrated that ultrathin a-SiC MEAs have a stable electrochemical profile and can record single unit action potentials chronically. They maintain a high signal-to-noise ratio and preserve both fast-spiking and slow-spiking units. We have demonstrated that the astroglial response to the a-SiC arrays is modest and extends, depth dependently, to not more than 250-350 μm from the electrode sites. Neuronal cell body counts were not different between implant and sham controls at either superficial or deep depths. Amorphous silicon carbide may provide a platform for the development of robust devices for chronic neural recording and stimulation. Future work remains to adapt this technology for use in non-human primate studies and potentially for human applications. Other challenges include the development of robust interconnects to preserve electrical connectivity between recording sites and external connectors.

## Author Contribution

Conceptualization: J.R.A., J.O.U., S.F.C., J.J.P.

Methodology: J.R.A., E.N.J., P.H., S.S.P., J.O.U., N.G., A.G.H.R., S.F.C., J.J.P.

Investigation: J.R.A., J.O.U., E.N.J, P.H., Y.W., R.R., B.S., S.N., S.P., A.G.H.R.

Data Curation: J.R.A., E.N.J, P.H., J.H., Y.M., V.D., G.V., K.D., A.S., T.T.

Writing – original draft: J.R.A., E.N.J, P.H.

Writing – review and editing: J.R.A., E.N.J, P.H., J.O.U., A.G.H.R., S.F.C., J.J.P. Supervision: S.F.C., J.J.P.

Funding Acquisition: S.F.C., J.J.P.

## Acknowledgements

We thank Shishir Waghray and Arya Raju for their assistance with immunohistochemical analysis.

## Institutional Review Board Statement

The study was conducted according to the guidelines approved by the Institutional Animal Care and Use Committee of The University of Texas at Dallas (Protocol 18-13 approved 26 September, 2018).

## Supplementary Figure

**Supplementary Figure:**
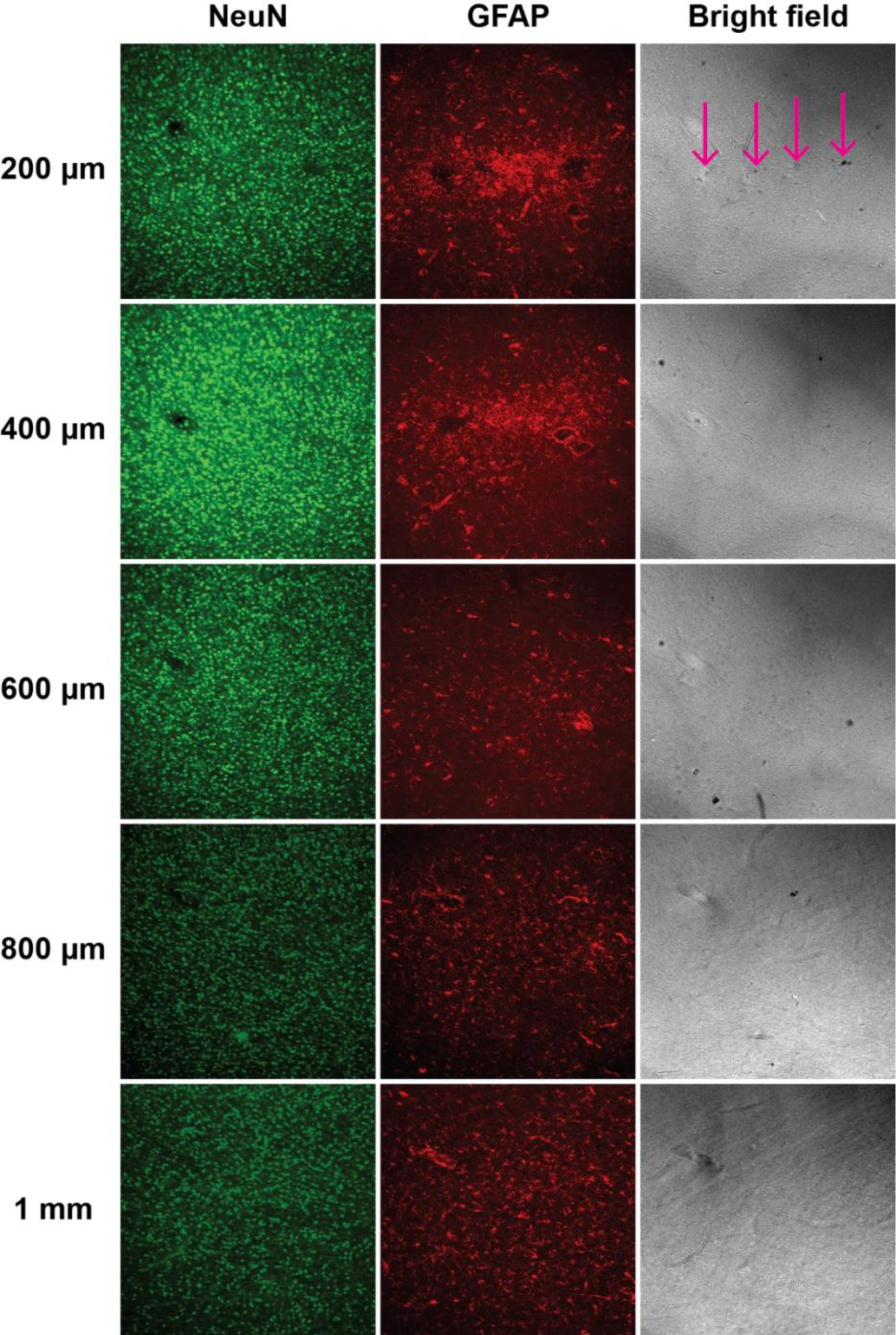
Slices from the same cortical column from 200 µm below the surface of the pia to 1 mm. The location of the implanted a-SiC probe becomes difficult to locate with increasing depth of the slices.

## References

[1] M. Velliste, S. Perel, M. C. Spalding, A. S. Whitford, and A. B. Schwartz, “Cortical control of a prosthetic arm for self-feeding,” Nature, vol. 453, no. 7198, pp. 1098–1101, Jun. 2008, doi: 10.1038/nature06996.

[2] S. N. Flesher et al., “A brain-computer interface that evokes tactile sensations improves robotic arm control,” Science, vol. 372, no. 6544, pp. 831–836, May 2021, doi: 10.1126/science.abd0380.

[3] V. L. Towle, T. Pham, M. McCaffrey, D. Allen, and P. R. Troyk, “Toward the development of a color visual prosthesis,” J. Neural Eng., vol. 18, no. 2, p. 023001, Apr. 2021, doi: 10.1088/1741-2552/abd520.

[4] T. W. Berger et al., “A Hippocampal Cognitive Prosthesis: Multi-Input, Multi-Output Nonlinear Modeling and VLSI Implementation,” IEEE Trans. Neural Syst. Rehabil. Eng., vol. 20, no. 2, pp. 198–211, Mar. 2012, doi: 10.1109/TNSRE.2012.2189133.

[5] M. L. Homer, A. V. Nurmikko, J. P. Donoghue, and L. R. Hochberg, “Sensors and Decoding for Intracortical Brain Computer Interfaces,” Annu. Rev. Biomed. Eng., vol. 15, no. 1, pp. 383–405, Jul. 2013, doi: 10.1146/annurev-bioeng-071910-124640.

[6] K. D. Harris and T. D. Mrsic-Flogel, “Cortical connectivity and sensory coding,” Nature, vol. 503, no. 7474, pp. 51–58, Nov. 2013, doi: 10.1038/nature12654.

[7] R. A. Normann and E. Fernandez, “Clinical applications of penetrating neural interfaces and Utah Electrode Array technologies,” J. Neural Eng., vol. 13, no. 6, p. 061003, Dec. 2016, doi: 10.1088/1741-2560/13/6/061003.

[8] D. R. Kipke, R. J. Vetter, J. C. Williams, and J. F. Hetke, “Silicon-substrate intracortical microelectrode arrays for long-term recording of neuronal spike activity in cerebral cortex,” IEEE Trans. Neural Syst. Rehabil. Eng., vol. 11, no. 2, pp. 151–155, Jun. 2003, doi: 10.1109/TNSRE.2003.814443.

[9] M. Armenta Salas et al., “Proprioceptive and cutaneous sensations in humans elicited by intracortical microstimulation,” eLife, vol. 7, p. e32904, Apr. 2018, doi: 10.7554/eLife.32904.

[10] C. E. Bouton et al., “Restoring cortical control of functional movement in a human with quadriplegia,” Nature, vol. 533, no. 7602, pp. 247–250, May 2016, doi: 10.1038/nature17435.

[11] S. N. Flesher et al., “Intracortical microstimulation of human somatosensory cortex,” Sci. Transl. Med., vol. 8, no. 361, Oct. 2016, doi: 10.1126/scitranslmed.aaf8083.

[12] T. D. Y. Kozai et al., “Comprehensive chronic laminar single-unit, multi-unit, and local field potential recording performance with planar single shank electrode arrays,” J. Neurosci. Methods, vol. 242, pp. 15–40, Mar. 2015, doi: 10.1016/j.jneumeth.2014.12.010.

[13] C. A. Chestek et al., “Long-term stability of neural prosthetic control signals from silicon cortical arrays in rhesus macaque motor cortex,” J. Neural Eng., vol. 8, no. 4, p. 045005, Aug. 2011, doi: 10.1088/1741-2560/8/4/045005.

[14] J. C. Barrese, J. Aceros, and J. P. Donoghue, “Scanning electron microscopy of chronically implanted intracortical microelectrode arrays in non-human primates.,” J Neural Eng, vol. 13, no. 2, Art. no. 2, 2016, doi: 10.1088/1741-2560/13/2/026003.

[15] P. R. Patel et al., “Utah array characterization and histological analysis of a multi-year implant in non-human primate motor and sensory cortices,” J. Neural Eng., vol. 20, no. 1, p. 014001, Feb. 2023, doi: 10.1088/1741-2552/acab86.

[16] J. M. Anderson, A. Rodriguez, and D. T. Chang, “Foreign body reaction to biomaterials,” Semin. Immunol., vol. 20, no. 2, pp. 86–100, Apr. 2008, doi: 10.1016/j.smim.2007.11.004.

[17] A. Lecomte, E. Descamps, and C. Bergaud, “A review on mechanical considerations for chronically-implanted neural probes,” pp. 0–12, 2017, doi: 10.1088/1361-6579/aa9828.

[18] J. N. Turner et al., “Cerebral Astrocyte Response to Micromachined Silicon Implants,” Exp. Neurol., vol. 156, no. 1, pp. 33–49, Mar. 1999, doi: 10.1006/exnr.1998.6983.

[19] V. S. Polikov, P. A. Tresco, and W. M. Reichert, “Response of brain tissue to chronically implanted neural electrodes,” J. Neurosci. Methods, vol. 148, no. 1, Art. no. 1, 2005, doi: 10.1016/j.jneumeth.2005.08.015.

[20] B. J. Black et al., “Chronic recording and electrochemical performance of Utah microelectrode arrays implanted in rat motor cortex,” J. Neurophysiol., p. jn.00181.2018, 2018, doi: 10.1152/jn.00181.2018.

[21] T. D. Y. Kozai, A. L. Vazquez, C. L. Weaver, S. G. Kim, and X. T. Cui, “In vivo two-photon microscopy reveals immediate microglial reaction to implantation of microelectrode through extension of processes,” J. Neural Eng., vol. 9, no. 6, Art. no. 6, 2012, doi: 10.1088/1741-2560/9/6/066001.

[22] S. M. Wellman et al., “Cuprizone-induced oligodendrocyte loss and demyelination impairs recording performance of chronically implanted neural interfaces,” Biomaterials, vol. 239, p. 119842, May 2020, doi: 10.1016/j.biomaterials.2020.119842.

[23] A. H. Marblestone et al., “Physical principles for scalable neural recording,” Front. Comput. Neurosci., vol. 7, 2013, doi: 10.3389/fncom.2013.00137.

[24] G. C. McConnell, H. D. Rees, A. I. Levey, C.-A. Gutekunst, R. E. Gross, and R. V. Bellamkonda, “Implanted neural electrodes cause chronic, local inflammation that is correlated with local neurodegeneration,” J. Neural Eng., vol. 6, no. 5, p. 056003, Oct. 2009, doi: 10.1088/1741-2560/6/5/056003.

[25] N. F. Nolta, M. B. Christensen, P. D. Crane, J. L. Skousen, and P. A. Tresco, “BBB leakage, astrogliosis, and tissue loss correlate with silicon microelectrode array recording performance,” Biomaterials, vol. 53, pp. 753–762, Jun. 2015, doi: 10.1016/j.biomaterials.2015.02.081.

[26] J. P. Seymour and D. R. Kipke, “Neural probe design for reduced tissue encapsulation in CNS,” Biomaterials, vol. 28, no. 25, pp. 3594–3607, Sep. 2007, doi: 10.1016/j.biomaterials.2007.03.024.

[27] J. G. Letner et al., “Post-explant profiling of subcellular-scale carbon fiber intracortical electrodes and surrounding neurons enables modeling of recorded electrophysiology,” J. Neural Eng., vol. 20, no. 2, p. 026019, Apr. 2023, doi: 10.1088/1741-2552/acbf78.

[28] T. D. Y. Kozai et al., “Ultrasmall implantable composite microelectrodes with bioactive surfaces for chronic neural interfaces,” Nat. Mater., vol. 11, no. 12, pp. 1065–1073, Dec. 2012, doi: 10.1038/nmat3468.

[29] L. Luan et al., “Ultraflexible nanoelectronic probes form reliable, glial scar–free neural integration,” Sci. Adv., vol. 3, no. 2, p. e1601966, Feb. 2017, doi: 10.1126/sciadv.1601966.

[30] D. Brassard and M. A. E. Khakani, “Dielectric properties of amorphous hydrogenated silicon carbide thin films grown by plasma-enhanced chemical vapor deposition,” J. Appl. Phys., vol. 93, no. 7, Art. no. 7, 2003, doi: 10.1063/1.1555676.

[31] G. L. Knaack, H. Charkhkar, S. F. Cogan, and J. J. Pancrazio, Amorphous SiliconCarbide for Neural Interface Applications. 2016. doi: 10.1016/B978-0-12-802993-0.00010-1.

[32] N. Geramifard, B. Dousti, C. Nguyen, J. Abbott, S. F. Cogan, and V. D. Varner, “Insertion mechanics of amorphous SiC ultra-micro scale neural probes,” J. Neural Eng., vol. 19, no. 2, p. 026033, Apr. 2022, doi: 10.1088/1741-2552/ac5bf4.

[33] S. F. Cogan, D. J. Edell, A. A. Guzelian, Y. P. Liu, and R. Edell, “Plasma-enhanced chemical vapor deposited silicon carbide as an implantable dielectric coating,” J. Biomed. Mater. Res. - Part A, vol. 67, no. 3, Art. no. 3, 2003, doi: 10.1002/jbm.a.10152.

[34] X. Lei et al., “SiC protective coating for photovoltaic retinal prosthesis,” J. Neural Eng., vol. 13, no. 4, p. 046016, Aug. 2016, doi: 10.1088/1741-2560/13/4/046016.

[35] U. Kalnins, A. Erglis, I. Dinne, I. Kumsars, and S. Jegere, “Clinical outcomes of silicon carbide coated stents in patients with coronary artery disease,” Med Sci Monit, vol. 8, no. 2, Art. no. 2, 2002.

[36] F. Deku, Y. Cohen, A. Joshi-Imre, A. Kanneganti, T. J. Gardner, and S. F. Cogan, “Amorphous silicon carbide ultramicroelectrode arrays for neural stimulation and recording,” J. Neural Eng., vol. 15, no. 1, Art. no. 1, 2018, doi: 10.1088/1741-2552/aa8f8b.

[37] E. N. Jeakle et al., “Chronic Stability of Local Field Potentials Using Amorphous Silicon Carbide Microelectrode Arrays Implanted in the Rat Motor Cortex,” Micromachines, vol. 14, no. 3, p. 680, Mar. 2023, doi: 10.3390/mi14030680.

[38] R. J. Vetter, J. C. Williams, J. F. Hetke, E. A. Nunamaker, and D. R. Kipke, “Chronic Neural Recording Using Silicon-Substrate Microelectrode Arrays Implanted in Cerebral Cortex,” IEEE Trans. Biomed. Eng., vol. 51, no. 6, pp. 896–904, Jun. 2004, doi: 10.1109/TBME.2004.826680.

[39] L. M. Frank, E. N. Brown, and M. A. Wilson, “A Comparison of the Firing Properties of Putative Excitatory and Inhibitory Neurons From CA1 and the Entorhinal Cortex,” J. Neurophysiol., vol. 86, no. 4, pp. 2029–2040, Oct. 2001, doi: 10.1152/jn.2001.86.4.2029.

[40] P. Stice, A. Gilletti, A. Panitch, and J. Muthuswamy, “Thin microelectrodes reduce GFAP expression in the implant site in rodent somatosensory cortex,” J. Neural Eng., vol. 4, no. 2, pp. 42–53, Jun. 2007, doi: 10.1088/1741-2560/4/2/005.

[41] J. Thelin et al., “Implant Size and Fixation Mode Strongly Influence Tissue Reactions in the CNS,” PLoS ONE, vol. 6, no. 1, p. e16267, Jan. 2011, doi: 10.1371/journal.pone.0016267.

[42] J. L. Skousen and P. A. Tresco, “The Biocompatibility of Intracortical Microelectrode Recording Arrays for Brain Machine Interfacing,” in Series on Bioengineering and Biomedical Engineering, 2nd ed., vol. 8, World Scientific, 2017, pp. 259–299. doi: 10.1142/9789813207158_0011.

[43] J. Maeng et al., “High-charge-capacity sputtered iridium oxide neural stimulation electrodes deposited using water vapor as a reactive plasma constituent,” J. Biomed. Mater. Res. B Appl. Biomater., vol. 108, no. 3, pp. 880–891, Apr. 2020, doi: 10.1002/jbm.b.34442.

[44] K. Krukiewicz, “Electrochemical impedance spectroscopy as a versatile tool for the characterization of neural tissue: A mini review,” Electrochem. Commun., vol. 116, p. 106742, Jul. 2020, doi: 10.1016/j.elecom.2020.106742.

[45] D. R. Merrill and P. A. Tresco, “Impedance Characterization of Microarray Recording Electrodes in Vitro,” IEEE Trans. Biomed. Eng., vol. 52, no. 11, pp. 1960–1965, Nov. 2005, doi: 10.1109/TBME.2005.856245.

[46] S. F. Cogan, “Neural Stimulation and Recording Electrodes,” Annu. Rev. Biomed. Eng., vol. 10, no. 1, Art. no. 1, 2008, doi: 10.1146/annurev.bioeng.10.061807.160518.

[47] J. D. Rolston, R. E. Gross, and S. M. Potter, “Common median referencing for improved action potential detection with multielectrode arrays,” in 2009 Annual International Conference of the IEEE Engineering in Medicine and Biology Society, Minneapolis, MN: IEEE, Sep. 2009, pp. 1604–1607. doi: 10.1109/IEMBS.2009.5333230.

[48] A. Stiller et al., “Mechanically Robust, Softening Shape Memory Polymer Probes for Intracortical Recording,” Micromachines, vol. 11, no. 6, p. 619, Jun. 2020, doi: 10.3390/mi11060619.

[49] J. O. Usoro et al., “Influence of Implantation Depth on the Performance of Intracortical Probe Recording Sites,” Micromachines, vol. 12, no. 10, p. 1158, Sep. 2021, doi: 10.3390/mi12101158.

[50] S. M. Wellman et al., “A Materials Roadmap to Functional Neural Interface Design,” Adv. Funct. Mater., vol. 28, no. 12, p. 1701269, Mar. 2018, doi: 10.1002/adfm.201701269.

[51] A. Joshi-Imre et al., “Chronic recording and electrochemical performance of amorphous silicon carbide-coated Utah electrode arrays implanted in rat motor cortex,” J. Neural Eng., vol. 16, no. 4, p. 046006, Aug. 2019, doi: 10.1088/1741-2552/ab1bc8.

[52] M. Okun, A. Lak, M. Carandini, and K. D. Harris, “Long Term Recordings with Immobile Silicon Probes in the Mouse Cortex,” PLOS ONE, vol. 11, no. 3, p. e0151180, Mar. 2016, doi: 10.1371/journal.pone.0151180.

[53] H. Zeng and J. R. Sanes, “Neuronal cell-type classification: challenges, opportunities and the path forward,” Nat. Rev. Neurosci., vol. 18, no. 9, pp. 530–546, Sep. 2017, doi: 10.1038/nrn.2017.85.

[54] S. P. Savya et al., “In vivo spatiotemporal dynamics of astrocyte reactivity following neural electrode implantation,” Biomaterials, vol. 289, p. 121784, Oct. 2022, doi: 10.1016/j.biomaterials.2022.121784.

[55] R. Lycke et al., “Low-threshold, high-resolution, chronically stable intracortical microstimulation by ultraflexible electrodes,” Cell Rep., vol. 42, no. 6, p. 112554, Jun. 2023, doi: 10.1016/j.celrep.2023.112554.

[56] A. Stiller et al., “A Meta-Analysis of Intracortical Device Stiffness and Its Correlation with Histological Outcomes,” Micromachines, vol. 9, no. 9, p. 443, Sep. 2018, doi: 10.3390/mi9090443.

[57] C. Beaulieu, “Numerical data on neocortical neurons in adult rat, with special reference to the GABA population,” Brain Res., vol. 609, no. 1–2, pp. 284–292, Apr. 1993, doi: 10.1016/0006-8993(93)90884-P.

